# Identification of conserved gene expression programs activated in multiple modes of torpor across vertebrate clades

**DOI:** 10.1101/2023.11.29.569284

**Authors:** Kurt Weir, Natasha Vega, Veronica F. Busa, Ben Sajdak, Les Kallestad, Dana Merriman, Krzysztof Palczewski, Joseph Carroll, Seth Blackshaw

## Abstract

Torpor encompasses adaptations to diverse extreme environmental stressors such as hibernation, aestivation, brumation, and daily torpor. Here we introduce StrokeofGenus, an analytic pipeline that identifies distinct transcriptomic states and conservation of gene expression patterns across studies, tissues, and species. We use StrokeofGenus to study multiple and diverse forms of torpor from publicly available RNA-seq datasets that span eight species and two classes. We identify three transcriptionally distinct states during the cycle of heterothermia: euthermia, torpor, and interbout arousal. We also identify torpor-specific gene expression patterns that are conserved both across tissues and between species with over three hundred million years of evolutionary divergence. We further demonstrate the general conservation of gene expression patterns in multiple forms of torpor, implying a common evolutionary origin for this process. Although here we apply StrokeofGenus to analysis of torpor, it can be used to interrogate any other complex physiological processes defined by transient transcriptomic states.

**HIGHLIGHTS:**

1. StrokeofGenus integrates orthologue annotation, non-negative matrix factorization, and transfer learning for cross-species analysis.
2. StrokeofGenus identifies conserved topor-related gene expression patterns across divergent species and cell types.
3. Heterothermia has three distinct transcriptomic states.
4. Torpor-specific gene expression patterns are conserved between species and forms of torpor.

## INTRODUCTION

To survive extreme ambient temperatures or food scarcity, some animal species dubbed heterotherms enter torpor, a state of inactivity accompanied by complex physiological changes. Torpid states are marked by suppressed heart rate, metabolic rate, body temperature, oxygen consumption, and blood flow as well as neuroprotective and muscle-protective mechanisms^1–3^, all of which contribute to large energy savings and resilience to muscle wasting and neurological and ischemic damage^2–10^.

Torpor is not limited to a single phylogenetic clade^1–3,5^. For instance, torpor strategies are used by a wide variety of mammals, birds, and reptiles^1–3,5^. Counterintuitively, heterotherms are often more closely related to non-heterotherms within the same clade than they are too other heterotherms^1–3^. It has therefore been suggested that torpor is an ancestral adaptation that has been repeatedly lost and that there are conserved regulatory mechanisms governing torpor between species^1–3^.

There are multiple named variations of torpor including hibernation, aestivation, daily torpor, and brumation. The criteria used to demarcate different forms is inconsistent, but generally include the length of the torpor bout, the season of inactivity, and/or taxonomic separation. For example, bouts of inactivity in daily torpor last less than a day while torpor bouts in hibernation can last weeks^2^. Aestivation is distinguished from hibernation by environmental temperature, with aestivation employed in exceedingly hot periods and hibernation in exceedingly cold conditions^11^. Like hibernation, brumation involves long torpor bouts during the cold season, but the term is applied specifically to reptilian species^5,8,9^. While these terms are segregated based on these criteria, no definitive answer has been provided regarding how mechanistically similar or dissimilar these torpor strategies are.

Hibernation is the most extensively studied form of torpor. It is characterized by long periods of very low activity and body temperature interrupted by periodic spikes to euthermic body temperature called interbout arousals (IBA). During this transition from torpor into IBA, gene expression is modified and global transcription as well as energy demands increase^4,10^. Despite the time points demonstrating similar body temperatures, IBA transcriptomes are distinct from euthermic time points^10^. Most early torpor transcriptomic studies have focused on the gene expression differences between euthermia and torpor^9,12–14^. However, due to the variability of activity between stages of torpor, more recent studies include multiple crucially timed samples throughout torpor, including pre- and post-torpor, IBA, and pre- and post-IBA time points^4,5,7,10,15^. Sampling many time points increases the complexity of analyzing gene expression changes using pairwise comparisons. Various groups have used different approaches to overcome this issue, such as setting the euthermic time point as a single point of comparison or adopting something like a figure-8 approach^4,10^. Often, time points are sampled and sequenced before it is known whether they represent a distinct transcriptomic state, further convoluting comparisons. There is a need for a technique that identifies distinct transcriptomic states and state-specific gene expression programs to simplify and clarify the calculation of stage-specific gene expression.

Independent transcriptomic-based studies have shown consistent gene expression changes during torpor within tissues across species. For instance, genes in the liver involved in carbohydrate catabolism and fatty acid synthesis were broadly downregulated while those related to fatty acid catabolism were broadly upregulated during torpor in the liver of the Chinese alligator (*Alligator sinensis*), Himalayan marmot (*Marmota himalayana*), and grizzly bear (*Ursus arctos horribilis*)^6–8^. Transcriptomic shifts involved in neuroprotection and protection against skeletal muscle atrophy have also been observed across species^4–6^. Recent work has used gene orthology to demonstrate shared molecular mechanisms during torpor among four mammalian species: the arctic ground squirrel (*Urocitellus parryii*), the 13-lined ground squirrel (13LGS, *Ictidomys tridecemlineatus*), the American black bear (*Ursus americanus*), and the Brandt’s bat (*Myotis brandtii)*^1^. However, there have to date been no reported comparisons between torpor transcriptomes in mammal and non-mammal species. Furthermore, comparisons across datasets based on the overlap of lists of significantly differential genes lack the capacity to demonstrate the conservation of global gene expression, which will involve many sub-significant but functionally related changes in gene expression.

We introduce a novel computational pipeline, dubbed StrokeofGenus, to publicly available torpor bulk RNA-seq datasets. Like prior pipelines, StrokeofGenus identifies orthology across species without the need for a reference genome to enable comparison of gene expression in non-model organisms^16^, but extends its functionality by including non-negative matrix factorization and transfer learning analysis for the discovery of complex patterns of co-expressed genes and the comparison of these patterns across datasets^17,18^. StrokeofGenus is able to identify transcriptomically distinct (and indistinct) phases of torpor and cross-study conservation of torpor gene expression programs. By identifying conserved gene expression programs across species, we demonstrate the presence of shared molecular mechanisms directing various forms of torpor, supporting the hypothesis that torpor is a shared ancestral trait.

## METHODS

### Sample Collection

Eleven female and four male 13-lined ground squirrels (13LGS) were obtained from the University of Wisconsin Oshkosh Squirrel Colony for use in this study. Animals were euthanized by decapitation under isoflurane anesthesia (isoflurane anesthesia was not used for winter torpor decapitations). The frontal cortex, hypothalamus, retina, RPE, and liver were dissected from each of three animals for five physiological states: summer euthermia, prehibernation/room temperature torpor, winter torpor, 3-days-post-arousal euthermia, and 14-days-post-arousal euthermia. Bio Medic Data Systems microchips and/or a FLIR thermal camera pointed at the inner ear were used to determine body temperature and combined with distinct behavioral phenotypes associated with torpor to determine physiological state. The experimental procedures described were approved by the Institutional Animal Care and Use Committee of the Medical College of Wisconsin (AUA00005654) and were in accordance with the ARVO Statement for the Use of Animals in Ophthalmic and Vision Research.

RNAse-free tubes and pipette tips were used. RNAse Zap was used on surfaces, dissection tools, and gloves before and between dissections. Sterile surgery personal protective equipment and tools were used. Tissues were transported in 1ml of RNAlater per 100mg of tissue.

### RNA sequencing

Tissue samples were removed from RNAlater, and total RNA was isolated with the miRNeasy micro kit with an optional DNase step, per the manufacturer’s protocol (Qiagen, Hilden, Germany). The total RNA was used to generate cDNA libraries with the Illumina TruSeq stranded Total RNA kit and sequenced on an Illumina Nextseq 500.

### *De novo* transcriptome generation, gene expression, and proteome inference

The bulk RNA-seq fastq files for each species were input to Trinity v2.13.1^19^ for *de novo* transcriptome assembly using default parameters. Every sample was used for each species, except for 13LGS, where, to save computation time with the high number and redundancy of samples, approximately half of the biological replicates were used (Supplementary Table 3). Gene expression for each sample was calculated using RSEM v1.2.15^20^ and bowtie2 v2.4.1^21^ implemented through the *align_and_estimate_abundance.pl* Trinity script. The *de novo* transcriptome fasta file output by Trinity was used as input for TransDecoder-v5.5.0^22^ to generate predicted proteomes with the “single_best_only” argument to produce only one protein sequence for each transcript sequence in the fasta file.

### Orthologue identification

Gene orthology relationships between species were reconstructed from the TransDecoder predicted proteome fasta outputs using OMAStandalone v2.5.0^23^ with the “DoHierarchicalGroups” parameter set to “false” and the “UseOnlyOneSplicingVariant” parameter set to “true” to generate gene-level relationships. Splice files, which are used by OMAStandalone to keep track of splicing variants of the same gene, were generated from the gene_trans_map output of Trinity using a custom script. The yeast proteome from the Orthology Matrix website was used as the out-group in orthology reconstruction in OMAStandalone^24,25^. Mouse, human, *D. melanogaster*, and *C. elegans* proteomes were downloaded from the Orthology Matrix website and included in orthology reconstruction as references for the quality of the *de novo* transcriptomes generated in this study.

### Pattern identification and conservation

Gene expression patterns were identified from the RSEM output for each dataset by nonnegative matrix factorization using the R package CoGAPS v3.10.0^17^. To reduce computation time, we used the *GWCoGAPS* function with the parameter *nSets* = 24 to split pattern finding across twenty-four groups of genes. Pattern markers were calculated for each pattern using the *patternMarkers* function. Conservation of gene expression patterns across datasets was calculated using the R package ProjectR v1.6.0^18^ with default parameters. Orthology information across species was imported from the OMA Standalone outputs OrthologousMatrix.txt and Map-SeqNum-ID.txt. Only genes with orthologs in both the reference and target datasets were considered.

### Prunetree diagram

The prunetree diagram (Fig S4A) was generated using species names on the TimeTree website^26^.

### Gene set enrichment analysis

Biological process gene ontology (GO) terms were downloaded from MSigD/B^27,28^. For each dataset, all gene names were converted to ortholog human gene symbols and ordered by CoGAPS pattern weights, which are non-negative. We then used the weighted approach implemented in GSEA^29^ via the R package fgsea v1.16.0^30^ with parameter *scoreType* = “pos” to test for enrichment of gene sets.

### Data Analysis

All data were processed and visualized using R version 4.0.2. Code to recapitulate all analyses is located at https://github.com/vbusa1/StrokeofGenus_manuscript.

## RESULTS

### StrokeofGenus enables cross-comparison of bulk RNA-seq torpor gene expression patterns across non-model species

We established an analysis pipeline able to reconstruct patterns of gene expression within bulk RNA-seq datasets, identify gene orthology across *de novo* transcriptomes, and compare torpor gene expression patterns across non-traditional model organisms (Fig 1A). First, bulk RNA-seq datasets for each species were fed into the *de novo* transcriptome assembly program Trinity^19^. From here, the pipeline bifurcates. Along one track, we calculated gene expression for each sample using Trinity’s native RSEM capability and the *de novo* transcriptome^20^. Gene expression information for each species was then fed to the non-negative matrix factorization R package CoGAPS^17^ to reveal the underlying structure of the data, including time point and tissue-specific patterns of gene expression. Along the other track, to obtain ortholog information across species, the *de novo* transcriptome was translated using Transdecoder^22^, and each species’ predicted proteome was fed into the orthology program OMA standalone^23^. OMA standalone identified the most similar orthologs for every gene across the species in question. Last, we input the OMA orthology information and CoGAPS-identified gene expression patterns into the transfer learning R package ProjectR^18^ to quantify the presence and strength of gene expression patterns across species.

**Figure 1:**
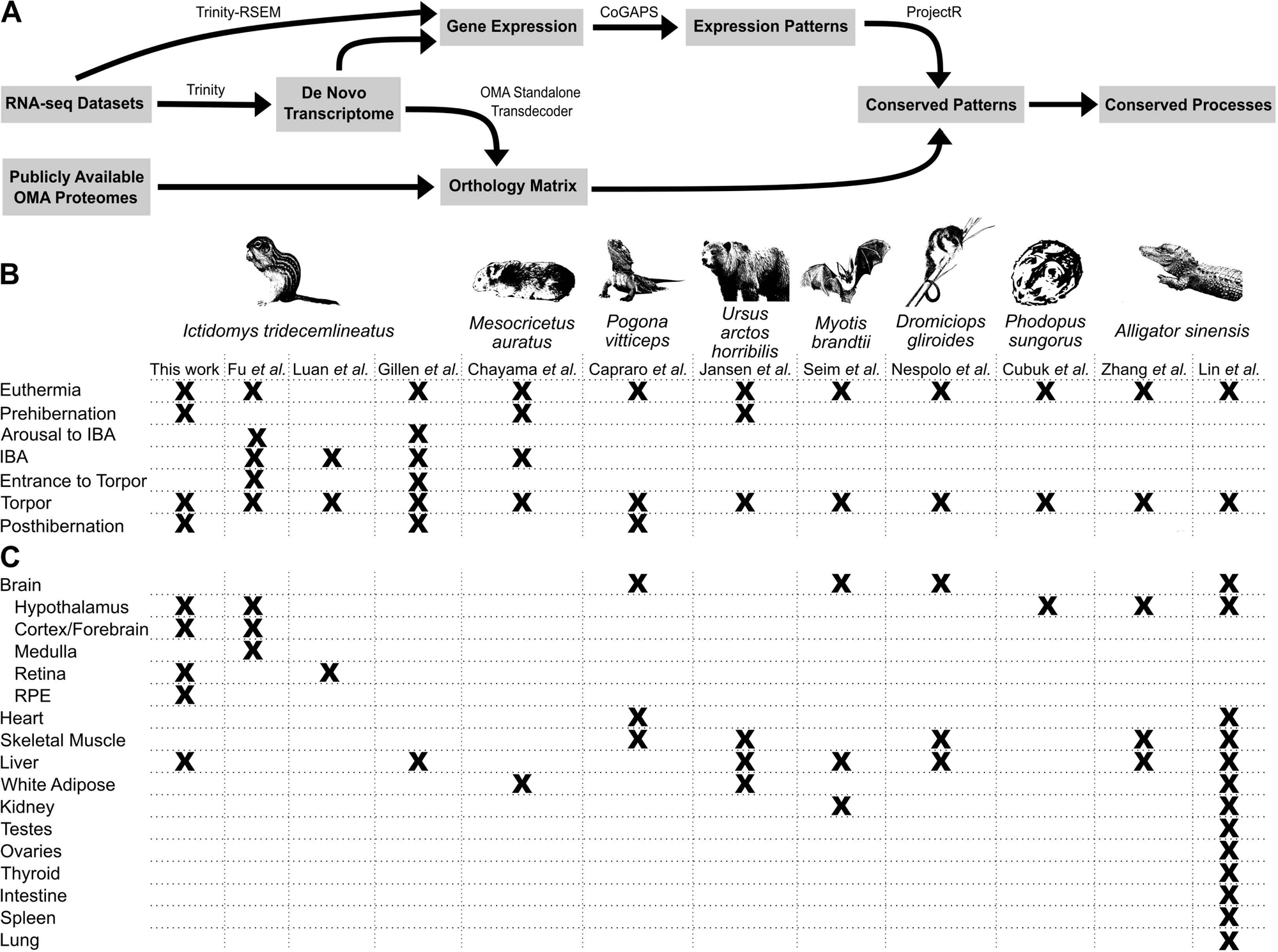
Overview of project design. (A) Flowchart for a novel analysis pipeline to reconstruct gene expression patterns and identify conservation of gene expression across species. The key software used in each step of the analytic pipeline is listed above each arrow. Table of datasets included in the study showing (B) time points and (C) tissues contained in each dataset.

There are publicly-available bulk RNA-seq datasets studying different forms of dormancy in species ranging from 13LGS to sea cucumber (*Apostichopus japonicus*) to Asiatic toads (*Bufo gargarizans*)^31–33^. To identify conserved patterns of gene expression throughout hibernation within vertebrate species, we selected a shortlist of eleven datasets in eight species with overlapping time points and tissues^4,5,7–10,12–15,31^ (Figure 1B, C). The datasets represent both mammals and reptiles that undergo variations of torpor including hibernation, daily torpor, brumation, and aestivation. The sampled tissues were enriched for central nervous system tissues and metabolically important tissues such as the liver. In addition, to further facilitate intra-species comparisons using StrokeofGenus, we generated a new 13LGS RNA-seq dataset that includes tissues for which data are publicly available in three other 13LGS hibernation datasets ^4,10,31^.

Between 7,578 and 19,422 orthologous genes were identified between all species comparisons (Supplementary Table 1). The number of genes reconstructed by Trinity is driven by the number of sequencing reads provided to the program^34^. Accordingly, larger numbers of genes were reconstructed for species with more samples (Supplementary Table 1). Species with greater numbers of reconstructed genes also had more orthologs identified across comparisons than those with fewer reconstructed genes, although this also appears to be impacted by evolutionary distance (Supplementary Table 1).

### Nonnegative matrix factorization identifies tissue- and time point-specific patterns of gene expression

To simultaneously uncover the distinct transcriptomic states and state-specific gene expression of the publicly available hibernation RNA-seq datasets, StrokeofGenus applies the matrix factorization tool CoGAPS. For almost all the datasets analyzed, CoGAPS was able to identify tissue and/or time point-specific patterns of gene expression (Fig 2, S1). CoGAPS was also able to deconvolve which time points within each dataset represented distinct transcriptomic states. We directed CoGAPS to identify between two and ten patterns for each dataset’s gene expression matrix. The results for each number of patterns were visualized as a heatmap showing the enrichment for each pattern in each sample. At low pattern numbers, samples were separated by tissue (Fig S2A). As the number of patterns increased, time point and tissue/time point-specific patterns emerged (Fig 2A). To determine how many patterns to derive from a dataset, we set the cutoff for hibernation-related patterns as the maximum number for which each pattern showed tissue and/or time point specificity but not sample specificity. If the number of patterns increased beyond this point, samples from the same time point and tissue would separate into sample-specific patterns (Fig S2B). We reasoned that only patterns with greater signal than sample-specific gene expression noise were biologically meaningful and useful for the purposes of discovery.

**Figure 2:**
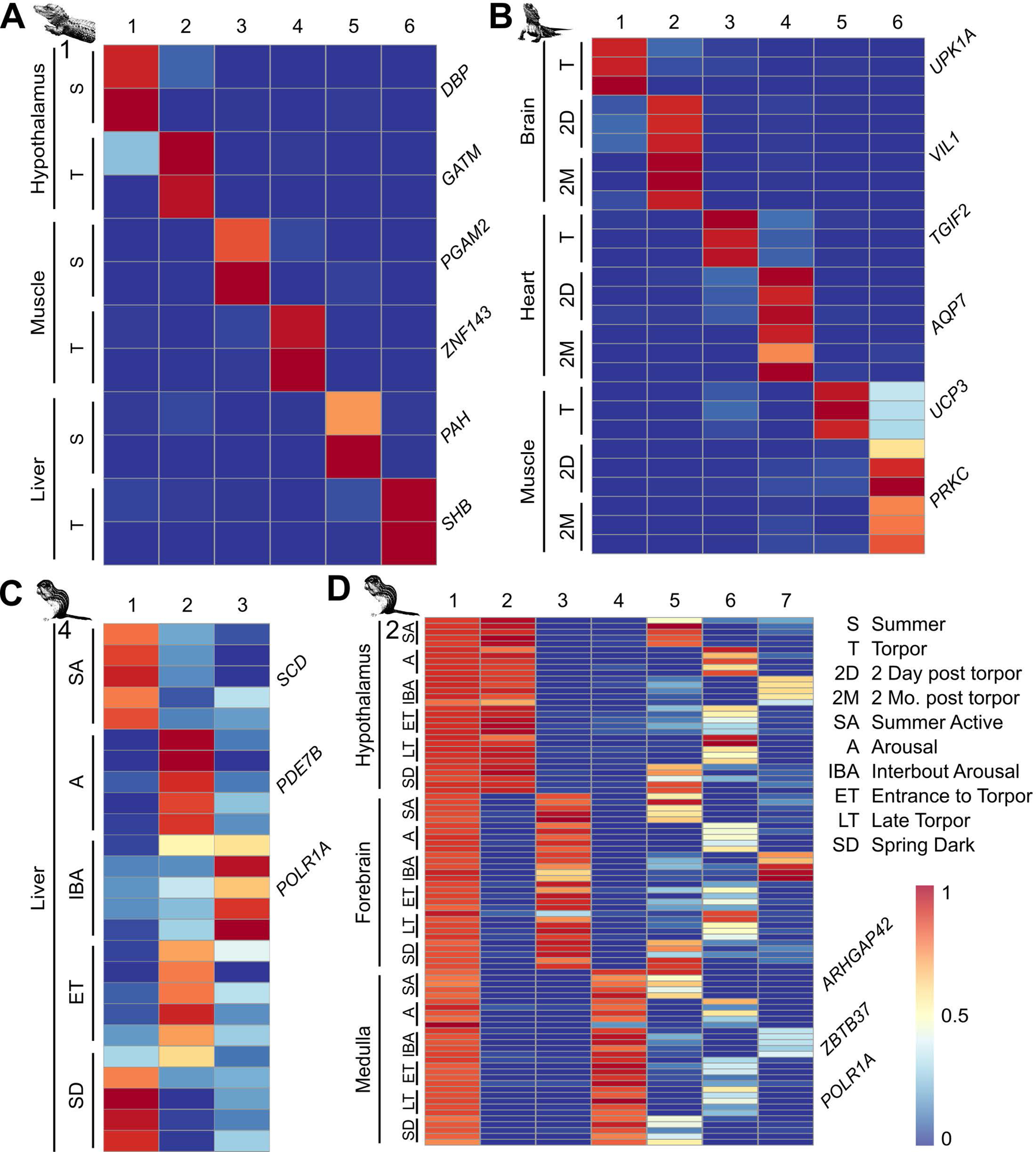
Identification of tissue and time point-specific gene expression patterns in torpor datasets. Heatmaps of tissue/time point-specific gene expression patterns in the (A) Chinese alligator dataset 1, (B) bearded dragon dataset, (C) 13LGS 4 dataset, and (C) 13LGS 2 dataset. Each row is a sample, and each column is a CoGAPS-derived pattern. Select pattern-specific genes that align with prior analyses are arrayed next to the relevant samples.

After optimizing the number of found patterns, we determined that some samples obtained from apparently distinct phases of hibernation represent equivalent transcriptomic states. For instance, the period immediately following exit from torpor was not transcriptomically distinct from a euthermic time point further temporally separated from torpor in the Australian bearded dragon (Fig 2B). We also found that the time points immediately preceding and following IBA are not distinct from deep torpor in 13LGS (Fig 2C). Our results corroborate the original publications’ findings, which were identified using principal component analysis (PCA) and random forest clustering techniques^4,5,10^, respectively. Based on our results, the principal transcriptomic states that characterize hibernation in mammals are euthermia, torpor, and IBA (Fig 2C, D).

Tissue and time point specificity manifested differently across datasets. In some datasets, individual patterns show combined tissue and time point specificity (Fig 2A, B). In other datasets, separate tissue-specific patterns were discovered along with distinct time point-specific patterns that were shared across tissues (Fig 2D, S2A). We observe that the latter case occurred in 13LGS 2, a dataset composed of closely related tissues, whereas the former occurred in datasets containing more distantly related tissues. This is likely because more closely related tissues share essentially the same gene expression changes while more distantly related tissues have distinct gene expression programs during torpor.

To identify genes with tissue- and state-specificity, we calculated the pattern marker genes for each pattern in each dataset (Supplementary Table 2, listed in order of pattern specificity). The top pattern-specific genes from our analysis aligned with those identified in prior analyses via pairwise comparisons, such as the circadian clock gene *DBP* upregulated in the Chinese alligator hypothalamus in summer^8^ and the transcriptional repressor *TGIF2* upregulated in the torpid heart of the bearded dragon^5^ (Fig 2).

### Transfer learning reveals the conservation of tissue and time point-specific gene expression across datasets

To determine whether there is conservation of torpor gene expression patterns across datasets, StrokeofGenus applies the transfer learning R package ProjectR^18^. ProjectR takes as input gene weights from a pattern learned in one dataset and tests for their enrichment in the samples of a second dataset. Samples whose gene expression profile matches the tested pattern receive higher ProjectR scores. If a pattern from a tissue and/or time point in one dataset shows enrichment in the equivalent samples of another dataset, this indicates that the datasets have shared gene expression programs. As proof of principle, we first applied ProjectR to different datasets generated from the same species with overlapping tissues. For this purpose, we used our 13LGS 1 dataset, which profiles multiple tissues, as a scaffold to compare with other 13LGS datasets that profile only one or a few tissues.

We first explored the conservation of the time point- and tissue-specific patterns of gene expression between the 13LGS 4 and 13LGS 1 datasets. Pattern 2 from the 13LGS 4 dataset is associated with torpid samples (Fig 2C). When projected into the 13LGS 1 dataset, this pattern showed greatest variance within liver samples and significant enrichment in late torpor liver samples relative to euthermic time points (Fig 3C, euthermia *P* = 3.2e-3, 2 days post hibernation *P* = 2.7e-3, 14 days post hibernation *P* = 1.1e-3, student’s t-test), consistent with a shared torpor-specific gene expression program. Gene Pattern 1 of the 13LGS 4 dataset demonstrated a distinct skew towards warm homeothermic states bracketing torpor (Fig 2C). Projection of pattern 1 into the 13LGS 1 dataset also demonstrated greatest variance in liver samples and significant separation of active liver samples from torpid samples (Fig 3C, euthermia *P* = 5.1e-6, 2 days post hibernation *P* = 2.8e-3, 14 days post hibernation *P* = 1.8e-3). Though ProjectR identified differences in gene expression between time points in the liver, CoGAPS analysis of the 13LGS 1 dataset only produced a general liver pattern 20 that did not identify any time point-specific gene expression (Fig S1A). This demonstrates that ProjectR can discern gene expression differences between samples when other sensitive methods are unable to do so. Interestingly, when pattern 20 of the 13LGS 1 dataset was projected into the 13LGS 4 dataset, it was enriched in active time points (Fig S3E, post hibernation/entrance *P* = 0.016).

**Figure 3:**
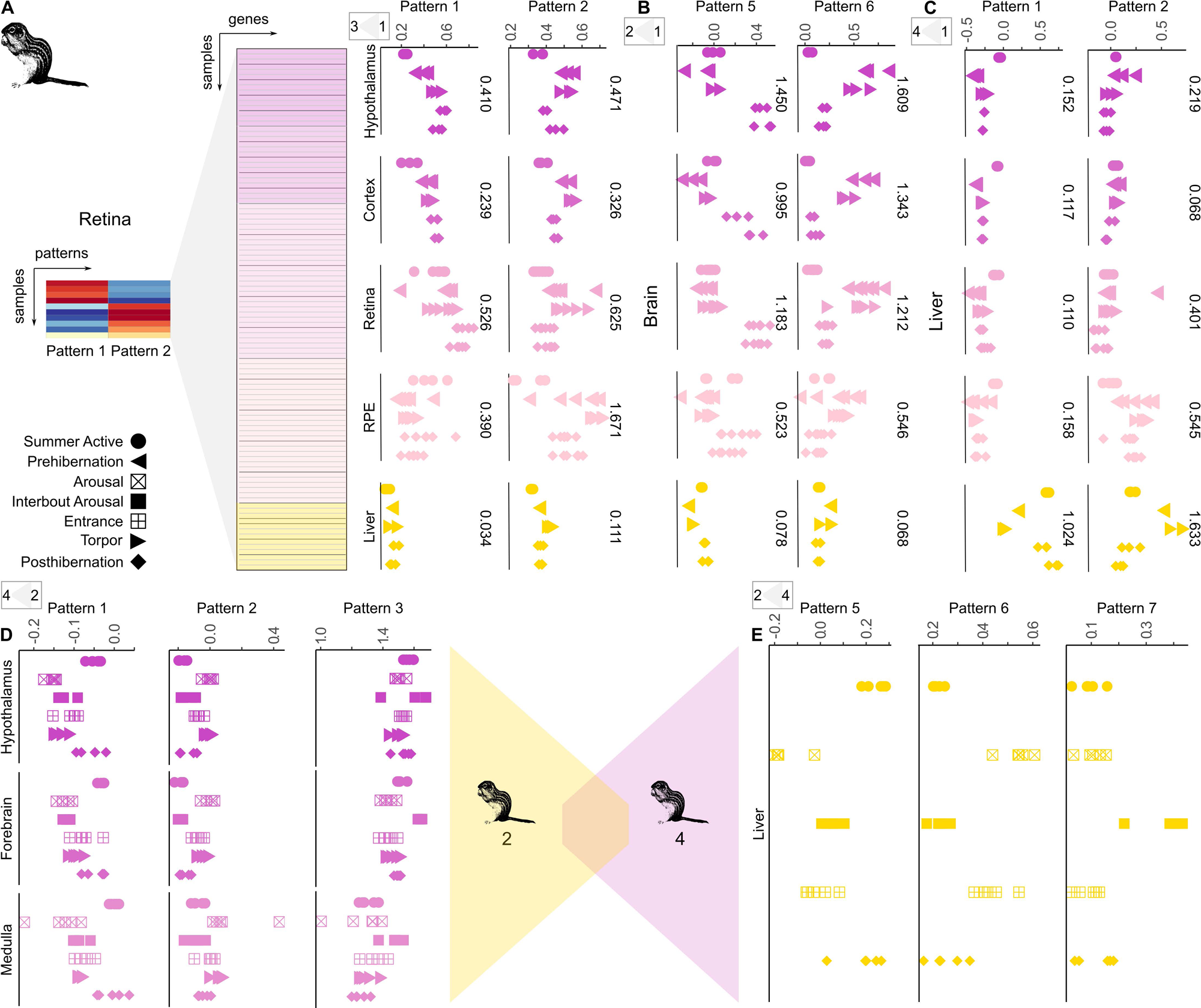
Conservation of gene expression patterns across datasets. Dot plots displaying tissue and time point-specific expression in the 13LGS 1 dataset of patterns from the (A) 13LGS 3, (B) 13LGS 2, and (C) 13LGS 4 datasets. (D) Dot plots displaying tissue and time point-specific expression in the 13LGS 2 dataset of patterns from the 13LGS 4 dataset. (E) Dot plots displaying time point-specific expression in the 13LGS 4 dataset of patterns from the 13LGS 2 dataset. Each point represents a single sample. RPE= retinal pigment epithelium

We further compared the 13LGS 2 dataset, which contains only brain tissue samples, with the 13LGS 1 dataset. Pattern 6 in the 13LGS 2 dataset, which is enriched in torpid samples (Fig 2D), showed greater variance within the four neural-related tissues than the liver samples in the 13LGS 1 dataset (Fig 3B). In the hypothalamus, Pattern 6 showed significant enrichment in torpid samples relative to active time points (euthermia *P* = 0.014, 2 days post hibernation *P* = 0.031, 14 days post hibernation *P* = 0.022). Similarly, Pattern 5, which is enriched in euthermic samples (Fig 2D), demonstrated greatest variance in neural-related tissues and showed significant segregation between euthermic time points following torpor and torpid time points in the 13LGS 1 hypothalamus sample (Fig 3B, 2 days post hibernation *P* = 5.1e-4, 14 days post hibernation *P* = 3.5e-3). Pattern 7 from the 13LGS 1 dataset, which was enriched in 14-day post hibernation hypothalamus samples (Fig S1A), was also enriched in euthermic samples in the hypothalamus of the 13LGS 2 dataset, while the torpid Pattern 5 (Fig S1A) showed enrichment in the torpid 13LGS 2 hypothalamus samples (Fig S3D, Pattern 7 euthermic/torpor p < 2.2e-16, Pattern 5 euthermic/torpor p < 2.2e-16).

The 13LGS 3, which includes only retinal tissue, and 13LGS 1 datasets also demonstrate conserved gene expression. Retinal euthermic Pattern 14 and the torpid Pattern 11 found in the 13LGS 1 dataset (Fig S1A), showed significant enrichment in the euthermic and torpid samples, respectively, in the 13LGS 3 dataset (Fig S3C, Pattern 14 *P* = 3.3e-4, Pattern 11 *P* = 4.6e-3). Similarly, torpid Pattern 2 from the 13LGS 3 dataset (Fig S1G) demonstrated the greatest variance in neural-related tissues and significant separation of torpid samples in the retina (Fig 3A, euthermia p = 2.5e-3, 2 days post hibernation *P* = 5.7e-3, 14 days post hibernation *P* = 7.7e-3), while the euthermic Pattern 1 showed enrichment in euthermic retinal samples immediately following torpor in 13LGS 1 (Fig 3A, 2 days post hibernation *P* = 3.7e-3, 14 days post hibernation p = 8.4e-3). These results cumulatively demonstrate that StrokeofGenus can identify conserved gene expression programs across multiple datasets generated from different laboratories.

To demonstrate that the functionality of StrokeofGenus is not limited to mammals, we additionally compared two datasets from the Chinese alligator and found conservation of euthermic and torpor gene expression for each tissue shared between the two datasets (hypothalamus, skeletal muscle, and liver), though the limited number of replicates prevents statistical comparisons (Fig S3A, B). These results further demonstrate the ability of ProjectR to identify conserved gene expression programs despite disparate collection times.

We also found conservation of gene expression patterns across tissues within the 13LGS. For example, the euthermic Pattern 5, torpid Pattern 6, and IBA Pattern 7 found in the 13LGS 2 (Fig 2D) demonstrate significant enrichment in the matching time points of the 13LGS 4 (Fig 3E, Pattern 5 euthermic/entrance *P* = 1.1e-4, Pattern 6 euthermic/entrance *P* = 1.3e-3, Pattern 7 IBA/entrance *P* = 6.2e-4). Similarly, projection of euthermic Pattern 1, torpid Pattern 2, and IBA Pattern 3 liver patterns into the 13LGS 2 dataset demonstrated coordinated upregulation in the corresponding time points in the forebrain (Fig 3D, Pattern 1 euthermic/torpor *P* = 1.1e-4, Pattern 2 euthermic/torpor *P* = 1.6e-4, Pattern 3 IBA/torpor *P* = 2.1e-4). Thus, within the same species, patterns of gene expression in hibernation are shared not only across datasets but also across tissues.

### Torpor gene expression programs are shared across species and different forms of torpor

Transfer learning provides an opportunity to quantify the conservation of torpor gene expression patterns across species and even across forms of torpor. Therefore, we used ProjectR in StrokeofGenus to test for gene expression conservation in the most common tissues across datasets–brain, liver, skeletal muscle, and white adipose– representing different taxonomic groups and forms of torpor.

To determine whether we could identify conservation of torpor gene expression patterns between species, we compared liver torpor expression between the grizzly bear and the 13LGS (most recent common ancestor (MRCA) ∼ 90 million years ago (MYA), Fig S4A), both of which hibernate. Pattern 3 from the grizzly dataset showed significant enrichment in torpid liver samples, whereas Pattern 2 is enriched in euthermic liver samples (Fig S1C). When projected into the 13LGS 4 dataset, these two grizzly patterns showed significant enrichment in the torpid and active 13LGS samples, respectively (Fig 4A, Pattern 3 euthermia/entrance *P* = 3.6e-3, Pattern 2 euthermia/entrance *P* = 1.8e-4). We performed the reciprocal comparison, projecting the 13LGS 4 dataset into the grizzly dataset, and found that Patterns 2, enriched in torpid time points, and Pattern 1, enriched in euthermic time points (Fig 2C), showed enrichment in the corresponding grizzly liver samples (Fig 4B, Pattern 2 euthermia/torpor *P* = 0.016, Pattern 1 euthermia/torpor *P* = 9.2e-4). Pattern markers from the source dataset with orthologs in the target dataset showed matching expression in the target dataset (Fig S4G).

**Figure 4:**
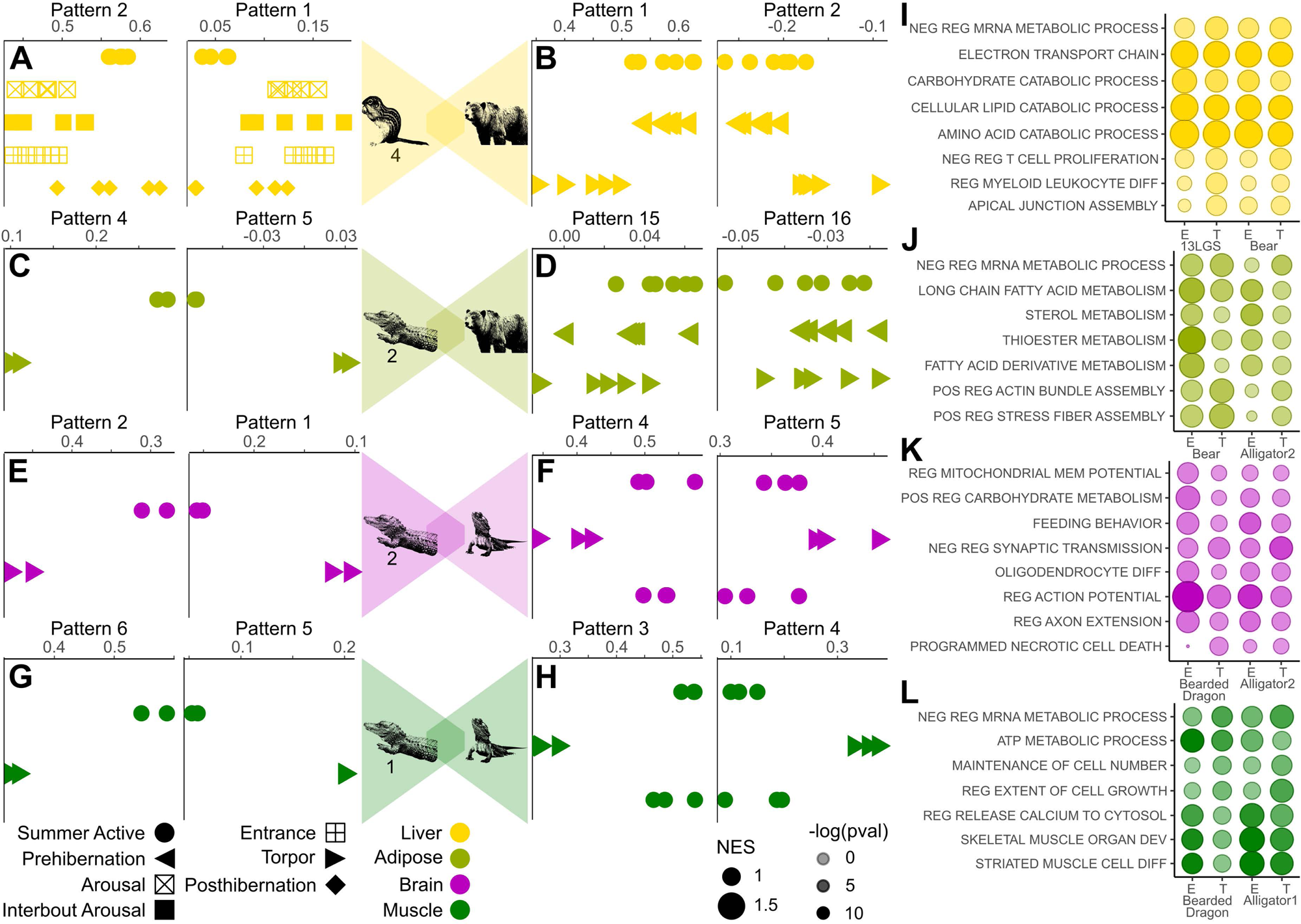
Conservation of gene expression patterns across species. (A) Dot plots displaying time point-specific expression in the 13LGS 4 dataset of torpor and euthermic patterns from liver samples in the grizzly dataset. (B) Dot plots displaying time point-specific expression in the liver samples of the grizzly dataset of torpor and euthermic patterns from the 13LGS 4 dataset. (C) Dot plots displaying time point-specific expression in the white adipose tissue (WAT) samples of the Chinese alligator 2 dataset of torpor and euthermic patterns from the WAT samples of the grizzly dataset. (D) Dot plots displaying time point-specific expression in the WAT samples of the grizzly dataset of torpor and euthermic patterns from the WAT samples of the Chinese alligator 2 dataset. (E) Dot plots displaying time point-specific expression in the hypothalamus samples of the Chinese alligator 2 dataset of torpor and euthermic patterns from the brain samples of the bearded dragon dataset. (F) Dot plots displaying time point-specific expression in the brain samples of the bearded dragon dataset of torpor and euthermic patterns from the hypothalamus samples of the Chinese alligator 2 dataset. (G) Dot plots displaying time point-specific expression in the skeletal muscle samples of the Chinese alligator 1 dataset of torpor and euthermic patterns from the skeletal muscle samples of the bearded dragon dataset. (H) Dot plots displaying time point-specific expression in the skeletal muscle samples of the bearded dragon dataset of torpor and euthermic patterns from the skeletal muscle samples of the Chinese alligator 1 dataset. Each point represents a single sample. Dot plots displaying the enrichment of biological process gene sets in torpor (T) and euthermic (E) patterns in (I) liver in 13LGS 3 and grizzly, (J) in adipose in Chinese alligator 2 and grizzly, (K) in hypothalamus/brain in Chinese alligator 2 and bearded dragon, and (L) in muscle in Chinese alligator 1 and bearded dragon. The X-axis corresponds to the CoGAPS pattern, and the Y-axis corresponds to the gene set. Dot size reflects the normalized effect size (NES) and the shade the -log(p-value) of enrichment.

We also found that Pattern 3 and Pattern 2 from the grizzly dataset were enriched in the equivalent time points in bat liver (MRCA ∼80 MYA, Fig S4A, C), though small sample numbers prevented statistical comparison, and Patterns 4 and 5 in the bat showed equivalent enrichment in the liver samples of the grizzly (Fig S4B, Pattern 4 euthermia/torpor *P* = 2.0e-8, Pattern 5 euthermia/torpor *P* = 0.014). This demonstrates the conservation of torpor gene expression across divergent mammalian species.

To identify whether the conservation of torpor gene expression is detectable over greater taxonomic distances, we compared white adipose tissue in the Chinese alligator 2 to the grizzly (MRCA ∼320 MYA, Fig S4A). When projected into the Chinese alligator 2 dataset, euthermic adipose Pattern 4 from the grizzly (Fig S1C) showed enrichment in euthermic adipose samples (Fig 4C). Similarly, projection of the torpid grizzly Pattern 5 (Fig S1C) into the Chinese alligator 2 dataset demonstrated segregation between torpid and euthermic replicates (Fig 4C), though small sample numbers prevented statistical comparisons in either case. Programs associated with euthermic adipose tissue in the Chinese alligator 2 dataset, Pattern 15 (Fig S1B), show enrichment in euthermic adipose tissue in grizzly (Fig 4D, euthermia/torpor *P* = 0.012). Taken together, these results suggest shared gene expression programs in torpor across vertebrate classes and hundreds of millions of years of evolutionary divergence.

To determine if distinct forms of torpor also share similar gene expression programs, we compared the Chinese alligator, which enters torpor in response to cold, to the bearded dragon, which enters torpor in response to extreme heat (MRCA ∼ roughly 280 MYA, Fig S4A). Euthermic and torpid brain patterns from bearded dragon showed enrichment in Chinese alligator 2 euthermic and torpid brain samples, respectively, though sample number precluded statistical comparisons (Fig 4E). Similarly, euthermic and torpid hypothalamus patterns 4 and 5 from the Chinese alligator 2 dataset significant enrichment in the euthermic and torpid samples of the bearded dragon brain, respectively (Fig 4F, Pattern 4 two days post torpor (2D)/torpor *P* = 0.017, two months post torpor (2M)/torpor *P* = 0.020, Pattern 5 2D/torpor, *P* = 0.048, 2M/torpor *P* = 0.082). Conservation of gene expression was also found between Chinese alligator brain patterns 2 and 3 and bearded dragon brain, though the separation between time points did not rise to statistical significance (Fig S4D). Conservation of torpor-specific gene expression between Chinese alligator and bearded dragon is also apparent in skeletal muscle (Fig 4G, H, Chinese alligator 1 dataset Pattern 3 2D/torpor *P* = 3.1e-3, 2M/torpor *P* = 1.2e-4, Pattern 4 2D/torpor *P* = 0.017, 2M/torpor *P* = 3.1e-4). We found greater conservation for patterns where conserved genes had greater pattern weight, meaning comparisons can be hampered over large taxonomic distances (Fig S4E, F). Not only is torpor gene expression shared across species and over large taxonomic distances, but across forms of torpor employed under opposite environmental conditions, such as extreme heat and cold.

To discern which biological processes comprise conserved gene expression programs, we applied gene set enrichment analysis (GSEA). We found enriched biological processes that agree with known phenotypes in torpor. Enrichment for gene sets was calculated for gene expression patterns within each species (Supplementary Table 4), then gene sets with similar enrichment were identified across species. As expected, mRNA production was downregulated in torpor across species and tissues^2^ (Fig 4I, J, L). Metabolic functions were upregulated in euthermic liver for both 13LGS and grizzly^2^ while immune cell development was selectively downregulated in torpid samples^35^ (Fig 4I). Adipose function was upregulated during euthermia in both grizzly and Chinese alligator 2^2^ (Fig 4J). As has been found in the Yakut ground squirrel brain^36^, actin fiber assembly was upregulated in both grizzly and Chinese alligator torpid adipose samples (Fig 4J). Like the results in liver and adipose, terms relating to tissue function were enriched in euthermic muscle in Chinese alligator 1 and bearded dragon (Fig 4L). Torpid animals retain muscle mass despite long periods of inactivity^2^ and in both Chinese alligator 1 and bearded dragon genes controlling maintenance of cell number were upregulated in torpid muscle samples (Fig 4L). Synapses retract during torpor before being rapidly regrown following a return to euthermia^3^. In both Chinese alligator 2 and bearded dragon brain samples, axon extension is upregulated in euthermic samples (Fig 4K). Interestingly, oligodendrocyte differentiation is also enriched in the euthermic brain in both Chinese alligator 2 and bearded dragon (Fig 4K). Demyelination of the hippocampus with the oligodendrocyte toxin cuprizone directs neurons to a dormant, axon-protective state in mice^37^. Not only do diverse species that employ different forms of torpor show conserved torpor gene expression programs, but these genes also regulate shared torpor-related biological processes.

## LIMITATIONS OF THE STUDY

Expression pattern conservation discoverability by StrokeofGenus is heavily dependent on study design. The determination of the correct number of patterns can be impacted when there are limited biological replicates. For instance, the Chinese alligator 2 dataset has only two replicates per condition and some sample-specific patterns were found before all condition-specific patterns were determined (Fig S1B). This limitation can be overcome by including a robust number of biological replicates (Fig 2, Fig S1). Also, StrokeofGenus has no function to directly determine which genes are driving the conservation of patterns across species. A strong inference can, however, be made by first identifying the global conservation of a pattern in the target dataset, and then identifying pattern markers whose expression in the target dataset aligns with that conservation (Fig S4E). This limitation could be eliminated in the future by the development of computational techniques to discern projection drivers. Overall, however, these limitations of StrokeofGenus are readily surmounted with robust and prudent study design.

## DISCUSSION

StrokeofGenus simplifies the analysis of time course gene expression data. Even with expanding numbers of time points, the identification of distinct transcriptomic states and state-specific gene expression is consolidated in a single step without the need for increasingly complex arrangements of pairwise comparisons. The matrix factorization output is also apt for cross-dataset and -species comparisons using transfer learning, enabling the identification of conserved gene expression without the need for manual comparison of gene lists.

With the combination of matrix factorization and transfer learning in StrokeofGenus, we identified the distinct transcriptomic states that compose different forms of torpor and demonstrated that some temporally distinct time points are transcriptomically identical. Distinct phases include euthermia, interbout arousal (IBA), and torpor (Fig 2, 3). In contrast, we show that pre- and post-hibernation and pre- and post-IBA are indistinguishable from euthermia and torpor, respectively (Fig 2, 3). Prior studies have suggested similar conclusions, but the broader view of the transcriptome considered via matrix factorization and transfer learning allows for clearer delineations between transcriptomic states.

Transfer learning demonstrates that torpor gene expression programs for various tissues such as the brain, liver, and white adipose tissue are shared across species, including between the mammal and reptile classes and between forms of torpor such as brumation and aestivation (Fig 4). This finding lends further support to the hypothesis that torpor is an ancestral adaptation that has been repeatedly lost rather than repeatedly independently evolved. Genes that drive conserved torpor patterns and which are found across species in this analysis are likely to be enriched for important torpor drivers and represent a pool of likely candidates for the manipulation of transcriptomic states. For instance, oligodendrocyte differentiation and axon extension show enrichment in the euthermic brain in both Chinese alligators and bearded dragons (Fig 4K). Genes directing these processes may play an important role in the brain’s rapid recovery and re-establishment of synapses following torpor^3^.

StrokeofGenus can be applied to further torpor and non-torpor questions. For instance, additional forms of torpor have been described in invertebrate species, such as aestivation in snails and sea cucumbers, the dauer state in nematodes, and diapause in insects^32,38–40^. Identification of conserved gene expression between torpor in more basal animals and those discussed in this paper could push back the date of the evolution of torpor and generalize the torpor state to include pauses in development. An additional goal of torpor research is to identify effective methods of inducing torpor in non-heterothermic species, which could improve organ transplantation and space flight^41–44^. Application of matrix factorization and transfer learning could determine how closely induced torpid states match naturally occurring torpor. Our pipeline could further be applied to identify conserved gene expression across species for shared processes other than torpor, such as shifting coat color in response to seasonal changes, post-infection immune recovery, or limb regeneration^45–47^. Any process that involves shifts between transcriptomic states and which is shared across species could be investigated for conserved gene expression using matrix factorization and transfer learning.

## DATA AVAILABILITY

Publicly available RNA-seq reads were downloaded from the European Nucleotide Archive for each species: 13LGS (PRJNA418486, PRJNA702062, PRJNA361561), Djungarian hamster (PRJNA743775), Australian central bearded dragon (PRJNA476034), grizzly bear (PRJNA413091), Brandt’s bat (SRP017183), monito del monte (PRJNA416414), Syrian hamster (PRJDB6278), and Chinese alligator (PRJNA593416, PRJNA556093). Raw sequencing reads generated for this study will be available upon publication on the Gene Expression Omnibus as GSE248932. All code used in the analysis is available at https://github.com/vbusa1/StrokeofGenus_manuscript.

## ACKNOWLEDGEMENTS

We thank the Johns Hopkins Transcriptomics and Deep Sequencing Core for assistance with generating RNA-Seq data and the lab of Elana Fertig for feedback on CoGAPS and ProjectR. This work was carried out at the Advanced Research Computing at Hopkins (ARCH) core facility (rockfish.jhu.edu), which is supported by the National Science Foundation (NSF) grant number OAC1920103. Research reported in this publication was supported by the National Eye Institute of the National Institutes of Health under award numbers R24EY027283 (to K.P. and S.B.), R21EY32281 (to S.B.), and F31EY031942 (to K.W.). The content is solely the responsibility of the authors and does not necessarily represent the official views of the National Institutes of Health. The authors acknowledge support to the Department of Ophthalmology Gavin Herbert Eye Institute at the University of California, Irvine from an unrestricted Research to Prevent Blindness award, and from NIH core grant P30EY034070.

## Supplemental Figures

**Figure S1:**
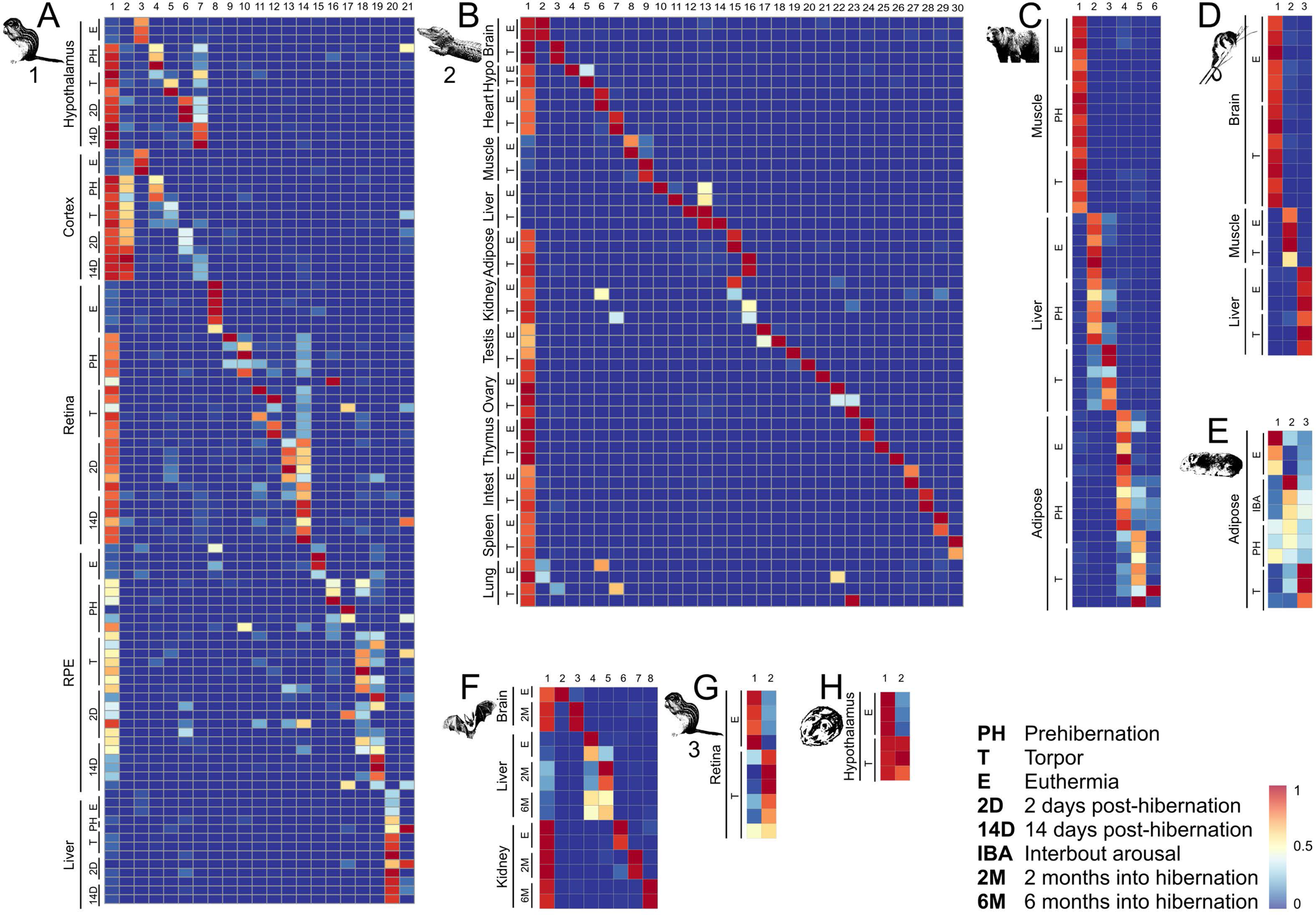
Identification of tissue and time point-specific gene expression patterns in torpor datasets. Heatmaps of tissue/time point-specific gene expression patterns in the (A) 13LGS 1 dataset, (B) Chinese alligator 2 dataset, (C) grizzly dataset, (D) monito del monte dataset, (E) Syrian hamster, (F) bat, (G) 13LGS 3, and (H) Djungarian hamster. Each row is a sample, and each column is a CoGAPS-derived pattern.

**Figure S2:**
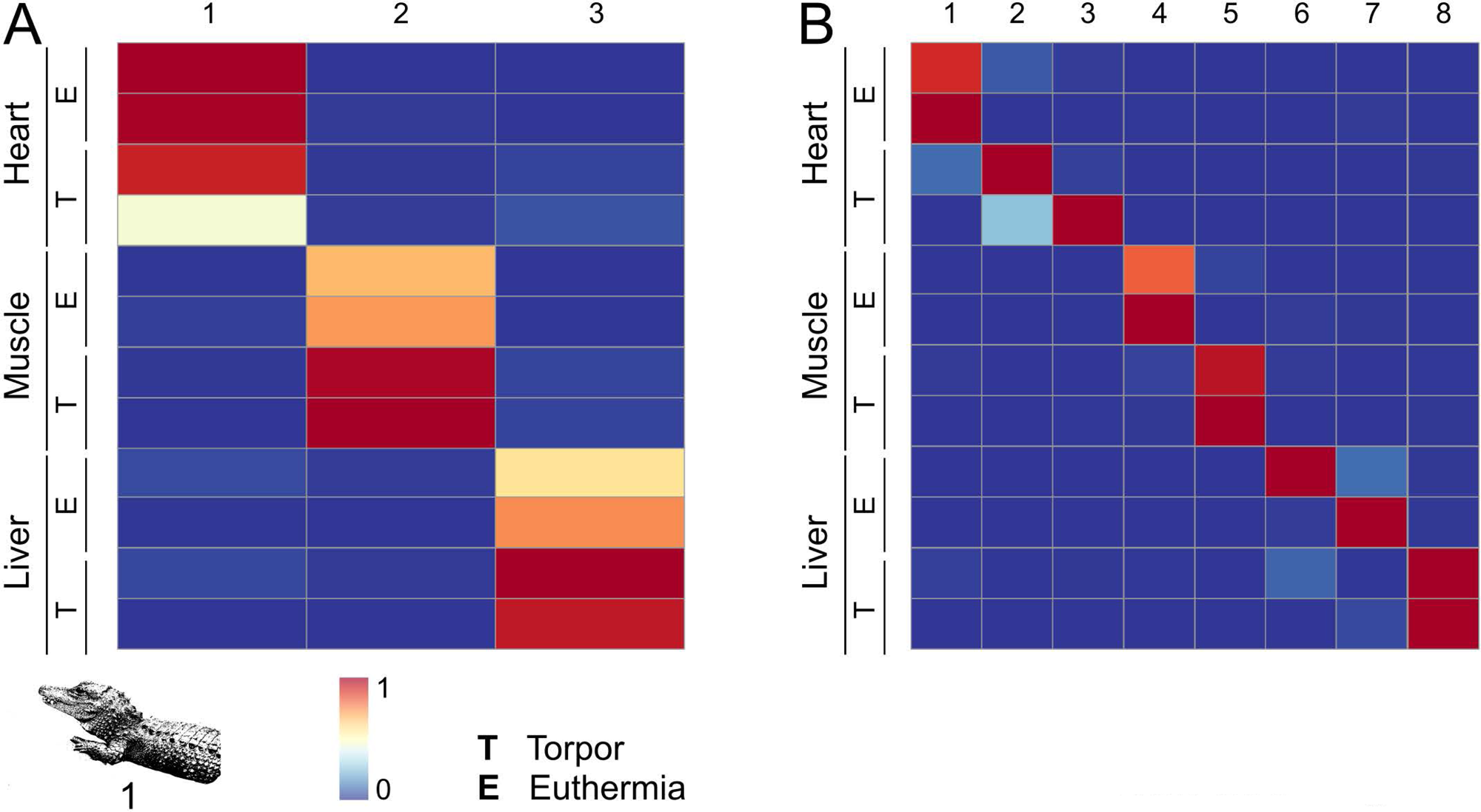
Strategy for determining pattern number in a dataset. Heatmaps of the Chinese alligator 1 dataset with (A) three patterns showing tissue-specific gene expression and (B) eight patterns showing either tissue and time point specificity or sample specificity. Each row is a sample and each column a CoGAPS-derived pattern.

**Figure S3:**
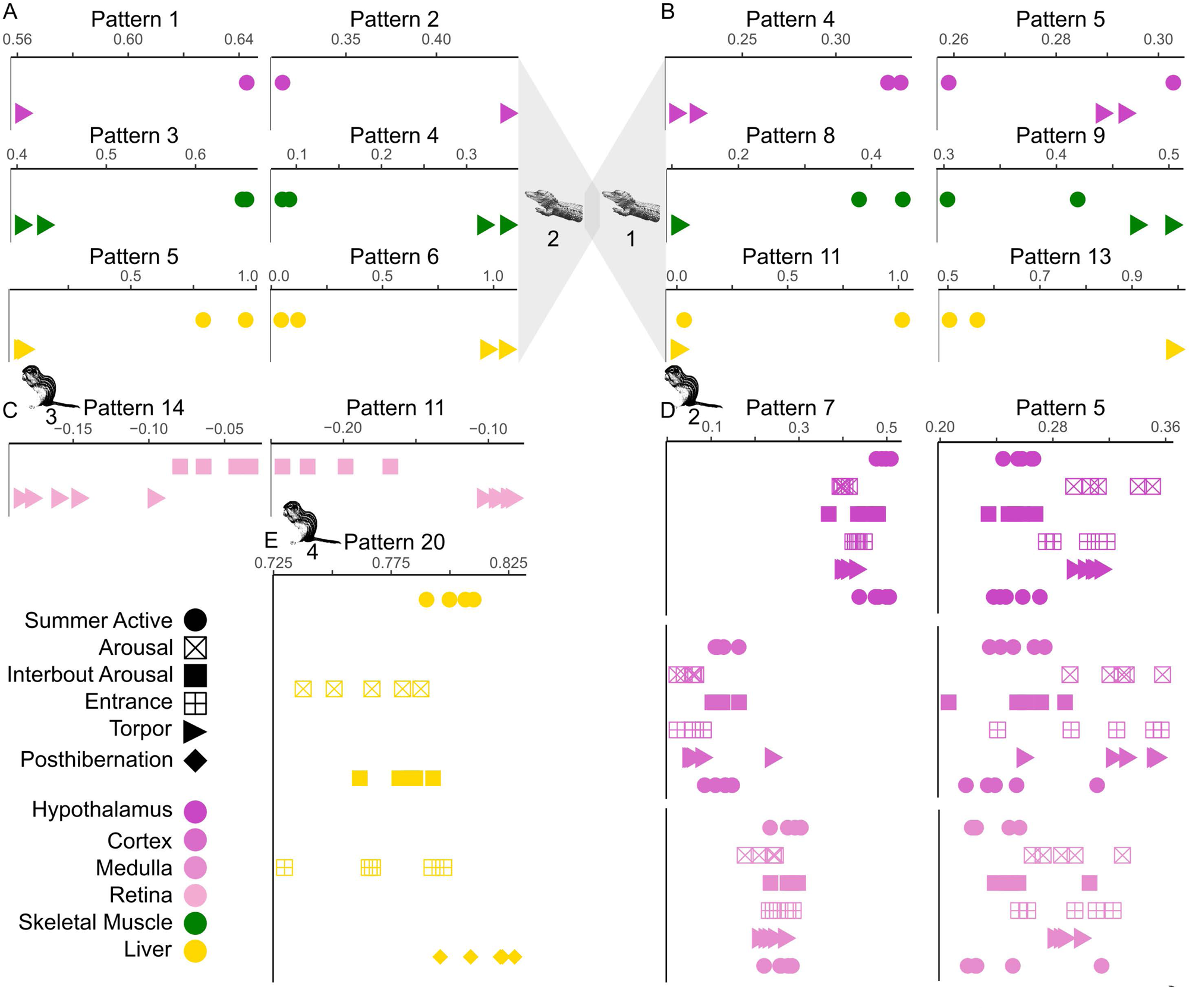
Conservation of gene expression patterns across datasets. Dot plots displaying time point-specific expression (A) in Chinese alligator 2 of patterns from Chinese alligator 1, (B) in Chinese alligator 1 of patterns from Chinese alligator 2, (C) in 13LGS 3 of patterns from 13LGS 1, (D) in 13LGS 2 of patterns from 13LGS 1, and (E) in 13LGS 4 of patterns from 13LGS 1. Each point represents a single sample.

**Figure S4:**
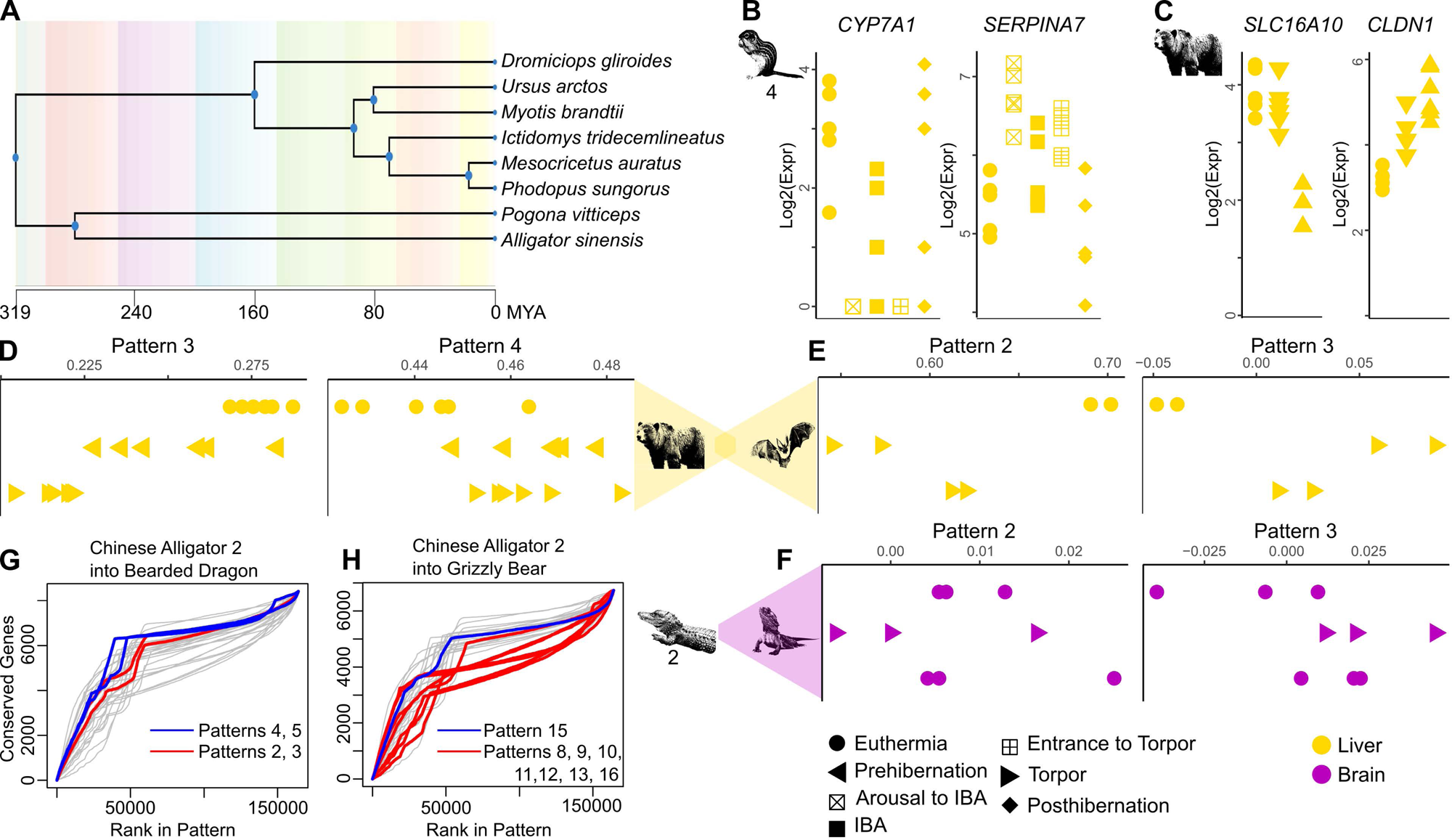
Conservation of gene expression patterns across species. (A) Dendrogram displaying hierarchical evolutionary relationships between species included in this study. Dot plots showing time point-specific expression of pattern markers from (B) grizzly in 13LGS 4 and (C) from 13LGS 4 in grizzly. Dot plots displaying time point-specific expression (D) in grizzly liver samples of patterns from the bat dataset (E) in bat liver samples of patterns from the grizzly dataset and (F) in bearded dragon brain samples of patterns from the Chinese alligator 2 dataset. Each point represents a single sample. Survival curves showing the rank for each pattern of the Chinese alligator 2 dataset for genes conserved with (G) bearded dragon and (H) grizzly. Blue lines show patterns with strong conservation across species and red lines show patterns with poor conservation. MYA = million years ago.

